# Robotic-inspired approach to multi-domain membrane receptor conformation space: theory and SARS-CoV-2 spike protein case study

**DOI:** 10.1101/2024.03.29.587391

**Authors:** Alen T. Mathew, Mateusz Sikora, Gerhard Hummer, A. Reza Mehdipour

## Abstract

The spike protein of SARS-CoV-2 is a highly flexible membrane receptor that triggers the translocation of the virus into cells by attaching to the human receptors. Like other type I membrane receptors, this protein has several extracellular domains connected by flexible hinges. The presence of these hinges results in high flexibility, which consequently results in challenges in defining the conformation of the protein. Here, We developed a new method to define the conformational space based on a few variables inspired by the robotic field’s methods to determine a robotic arm’s forward kinematics. Using newly performed atomistic molecular dynamics (MD) simulations and publicly available data, we found that the Denavit-Hartenberg (DH) parameters can reliably show the changes in the local conformation. Furthermore, the rotational and translational components of the homogenous transformation matrix constructed based on the DH parameters can identify the changes in the global conformation of the spike and also differentiate between the conformation with a similar position of the spike head, which other types of parameters, such as spherical coordinates, fail to distinguish between such conformations. Finally, the new method will be beneficial for looking at the conformational heterogeneity in all other type I membrane receptors.

## 2 Introduction

Membrane receptor proteins are found in all domains of life, making them a common feature across different organisms. They are essential in signaling, immune response, intercellular communication, and adhesion. Many of these receptors belong to the type-I transmembrane protein group [1]. They have a multi-domain extracellular N-terminus, followed by a single transmembrane helix and often a C-terminal intracellular domain. These receptors exhibit diverse architectural characteristics, which enable them to perform a wide range of functions. The extracellular part serves as a receiver or transmitter of signals, playing a crucial role in sensing and relaying communication between cells and their environment. A common feature of the extracellular parts is the presence of multiple domains connected by flexible hinges. This particular feature enables the extracellular part to be dynamic, which allows for specificity and accessibility outside of the cell. This also allows for large biologics such as antibodies to be accessed. However, these dynamic and flexible structures possess significant degrees of freedom, making it challenging to define their conformational state [2, 3]. The methods used to describe receptor conformation include measuring angles and distances between domains, determining the distance from a reference structure (X-ray or Cryo-EM), and using dimension reduction methods like principal component analysis to extract variables from internal coordinates. [4, 5]. These approaches provide valuable insights into receptor conformation, but they have limitations in defining intuitive collective variables for overall conformational changes. Therefore, there is a need for new methods for tracking conformational changes.

Kinematics is a branch of physics that deals with the description of motion. It focuses on the study of the motion of objects without considering the physical forces that cause the motion. When it comes to proteins, we want to determine which conformations are permissible while maintaining the protein’s structure without relying on forces to explain how the structure remains intact [6]. According to this definition, the analysis of motion (kinematics) is essentially a geometric problem. It is interesting to note that in the field of robotics, forward kinematics has been introduced to calculate the cartesian coordinates (x, y, z) of the robot’s end-effector or any of its intermediate joints using the robot’s internal coordinates (joint angles and link lengths). These approaches enable us to determine the configuration of the robotic arm by combining the conformation of its joints. This procedure is exciting to us since it embodies a set of variables that define the major degrees of freedom for the system based on the flexible joints (which are the hinges between the different domains of the protein).

The SARS-CoV-2 spike glycoprotein is also a type I membrane protein (Fig. 1a) and has 1273 amino acid residues, including an N-terminus signal peptide, a receptor-binding fragment S1 and a fusion fragment S2. S1 can be further divided into the N-terminal domain (NTD), receptor-binding domain (RBD), and C-terminal domains (CTD1 and CTD2). At the same time, S2 includes fusion peptide (FP), fusion-peptide proximal region (FPPR), heptad repeat 1 (HR1), central helix (CH), connector domain (CD), heptad repeat 2 (HR2), transmembrane segment (TM) and the cytoplasmic tail (CT) [7]. The spike protein is the receptor that the SARS-CoV-2 virus uses to attach to human cells via the ACE2 receptor, triggering viral entry [8]. Like other type-I receptors, the spike protein is dynamic due to several linked domains and flexible hinges. Several studies highlight the importance of spike flexibility for its function and the effect of glycans on its conformational heterogeneity [9, 10]. Therefore, defining the conformational space of the spike is highly desirable. However, like other receptors, several flexible hinges between domains complicate the definition of the variables accounting for the conformation space. Again, like other receptors, there have been some efforts to describe the conformational space of the spike protein. However, these studies have been mainly focused on a few angles and distances, which show the localized changes in the conformation and not the receptor’s conformation as a whole entity [7, 10, 11, 12].

**Figure 1.**
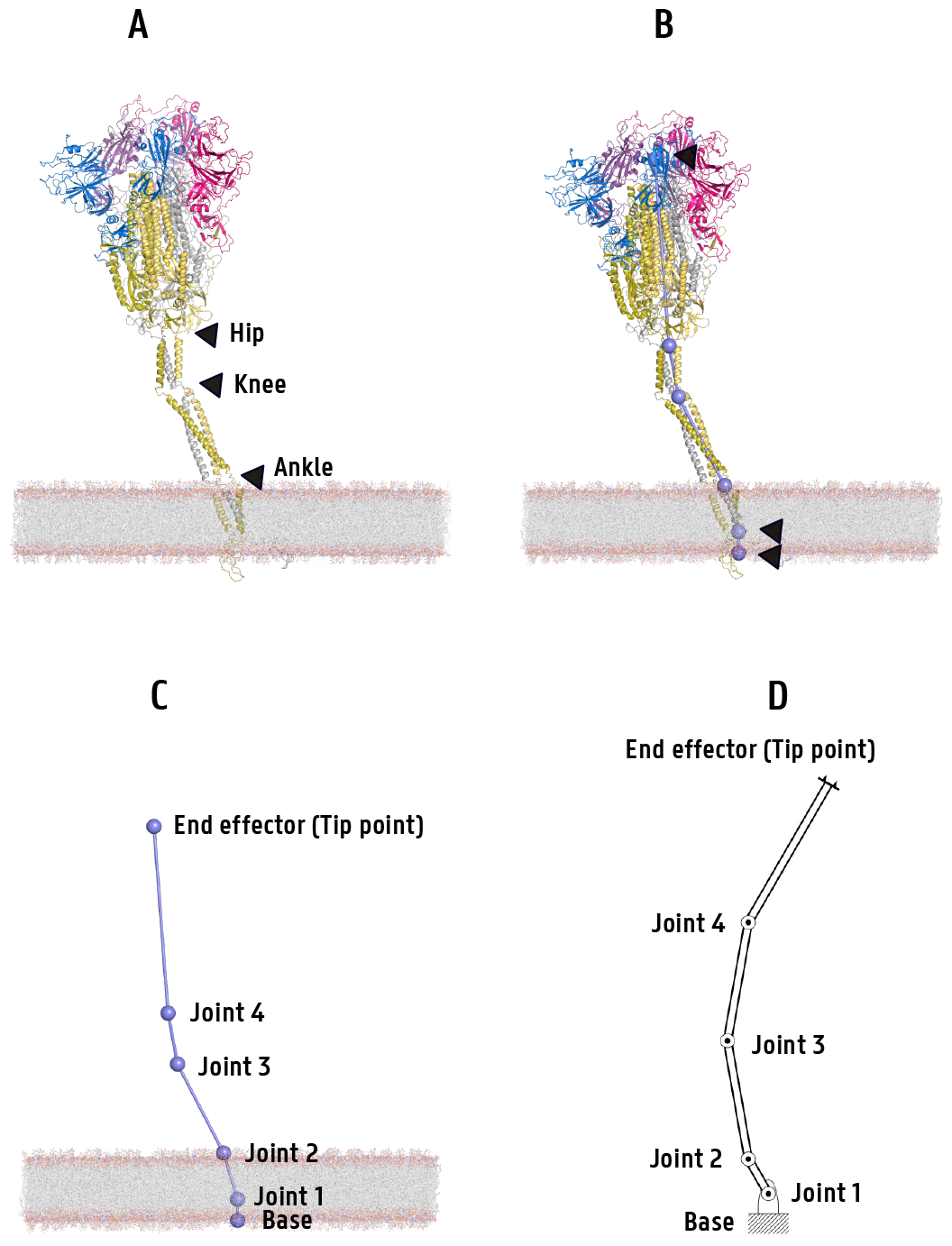
Spike as a robotic arm. **A** Previously defined hinges (joints) in the spike. **B** Three added points. **C** New of spike names for joints. **D** robotic-like scheme

This work aims to develop a mathematical framework based on the forward kinematic approaches used in robotics to define the space of receptor conformations. We introduce key concepts such as domains, and protein hinges as linkers and joints. Then, we apply geometric methods to precisely define the relationship between these coordinate sets and conformational space. Finally, our focus is on deriving a few collective variables that illustrate the conformational space of the protein as a robotic arm-like system.

## 3 Theory

### 3.1 Forward kinematics based on the Denavit-Hartenberg convention

We can assume that the spike is a robotic arm, as shown in Figure 1d. We opted to add three more points to the existing Ankle, Knee, and Hip defined joints. This will result in having five linkers and four joints, including the spike hinges and one additional joint inside the TM region (as seen in Fig. 1b, Fig. 1c, and Table S1). We can use the modified version of the definition of serial molecular chains proposed by Wright et al. [13] and Kavraki et al. [14]. The Denavit-Hartenberg (DH) parameters are commonly used to describe the transformation of the coordinate frame of each joint to the coordinate frame in the previous joint in the chains [15]. We used the DH convention to describe the forward kinematics of the robotic arm representation of the spike. For a robotic arm, we can define the DH parameters for each joint, which can be done for the spike protein (or, in general, membrane receptors). There are four joints, each with 3 degrees of freedom (DoF) and an additional translational DoF from the base to the first joint. Then, this robotic arm has 13 DoFs and represents the spike conformation. We can write a homogeneous transformation matrix (^*i*^*θ*_*i*+1_) for the frame joint i to transform to the frame of joint i-1 using three consecutive transformations (Eq. 1): (i) the translation along the (*Z*_*i*_) axis by *d*_*i*_, (ii) the rotation of (*Z*_*i*_) around the (*X*_*i*_) axis by *α*_*i*_, and (iii) the rotation of (*X*_*i*_) around the (*Z*_*i−*1_) axis by *θ*_*i*_ (Fig. S1).

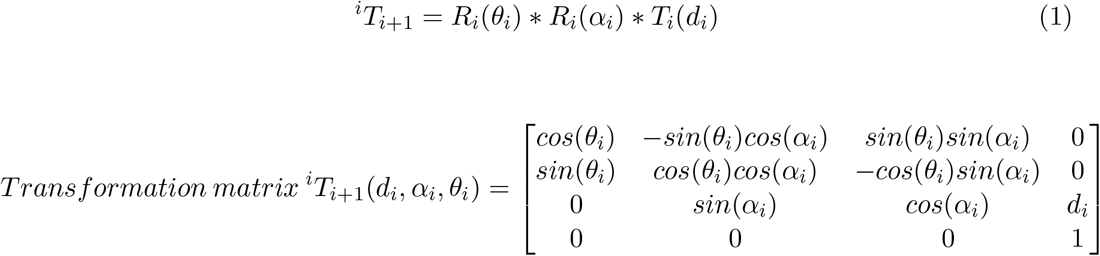

The only exception is the transformation from the base to the joint one (*A*_1_), a translation-only operation. Then, the transformation matrix of the whole chain to have the coordinates of the endpoint effector (tip of the S) in the base coordinate system will be:

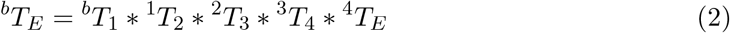

The elements of the final transformation matrix are listed in Appendix 1.

#### 3.1.1 Euler angles

Ultimately, we have a homogeneous 4*×*4 transformation matrix based on the DH parameters. It is a homogeneous matrix without scaling, the last row is always [0 0 0 1], and the number of actual variables is 4*×*3. Therefore, we must find a way(s) to reduce the dimension.

The homogeneous transformation matrix has two components: the rotation and translation parts:

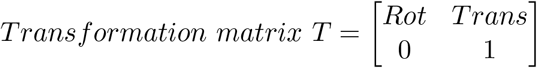

We calculated three Euler angles (*α,β,γ*) from the rotation component.

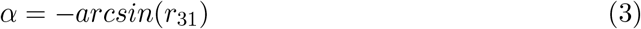

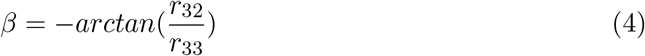

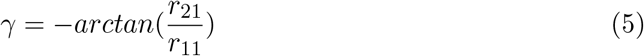

Also, from the translation part of the transformation matrix, we have three translational variables (*t*_1_ = *T*_*x*_, *t*_2_ = *T*_*y*_, *t*_3_ = *T*_*z*_)). At the end, there are six variables (*α, β, γ, T*_*x*_, *T*_*y*_, *T*_*z*_).

While the DH-based transformation matrix’s rotation part embodies the spike’s rotation information as a serial chain with four flexible joints, the translation part of the transformation matrix is equivalent to r, the distance between the base and the tip point.

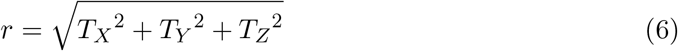

### 3.2 Forward kinematics based on the spherical coordinates (SC)

Another way of having forward kinematics is by looking at the tip point position and its position relative to the base point. Based on this approach, we can calculate the spherical parameters based on the vector between the base and tip points (Fig. S2). Here, we have three parameters: d is the vector (r) length, *θ*, and *φ* are spherical angles.

### 3.3. Distance metric

We can calculate the distance between two conformations using their transformation matrices by calculating the following left-invariant Riemannian distance metric [16]:

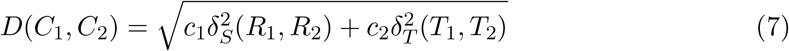

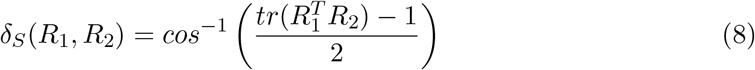

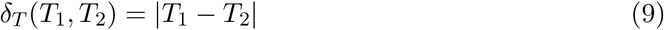

in which *δ*_*S*_ is the rotation Euler angle between the rotation components (*R*_1_,*R*_2_) of the transformation matrices of two conformations (*C*_1_,*C*_2_), and *δ*_*T*_ is the Euclidean distance between the translational components (*T*_1_,*T*_2_) of their transformation matrices. The parameters *c*_1_ and *c*_2_ are the weighting factors to homogenize the two terms since *δ*_*S*_ and *δ*_*T*_ have different measurement units (rad and nm, respectively). Here, we set *c*_1_ and *c*_2_ to 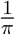 and 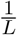 respectively. L is the maximum length of the translation part (about 30 nm).

For the SC parameters, to calculate the distance metric, we can build a single transformation matrix in which d is the length of the vector (r), *θ* is spherical angle *θ*, and *α* is spherical angle *φ*.

Then, the transformation matrix is:

#### Transformation matrix based on spherical parameters (^*b*^*T*_*E*_)

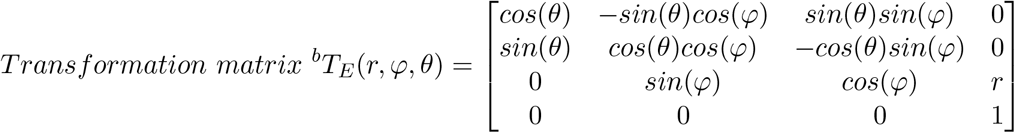

## 4 Results and Discussion

### 4.1 DH parameter distribution shows changes in both local and global conformations

To analyze the conformation of the spike protein, we performed triplicate simulations for both glycosylated and non-glycosylated pre-fusion spike proteins (3*×*500 ns amounted to 1.5 *µs* for each set of systems). We also analyzed simulations from 4 different labs using publicly available full-length spike protein simulation data (Tajkhorshid, Im, Klauda, and Hummer) (Table S2) [10, 12, 17, 18, 19, 20]. We analysed the 37.0 *µs* of the spike simulation data (31 *µs* glycosylated and 6.5 *µs* non-glycosylated simulations). Then, the DH parameters (5 linker lengths, 4 *α* angles, 4 *θ* angles) (Table 1 in appendix) were calculated for the simulations. Using equation (1), we calculated the transformation matrix for each joint. Then, using equation (2), we obtained the final transformation matrix. We also calculated the SC parameters (the distance r, two angles *θ* and *φ*).

**Table 1:** Denavit-Hartenberg parameters for spike joints transformation

| Transformation (${}^i\theta_{i+1}$) | $a_i$ | $d_i$ | $\alpha_i$ | $\theta_i$ |
| --- | --- | --- | --- | --- |
| ${}^bT_1$ | 0 | $d_0$ | 0 | 0 |
| ${}^1T_2$ | 0 | $d_1$ | $\alpha_1$ | $\theta_1$ |
| ${}^2T_3$ | 0 | $d_2$ | $\alpha_2$ | $\theta_2$ |
| ${}^3T_4$ | 0 | $d_3$ | $\alpha_3$ | $\theta_3$ |
| ${}^4T_E$ | 0 | $d_4$ | $\alpha_4$ | $\theta_4$ |

The distributions of DH parameters are shown in Fig. 2. As can be seen, The linker lengths also have narrow distributions. The distribution of DH *α* angles are restricted in the range betwen 0 and 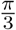. This is interesting because the *α* angles can theoretically sample values up to 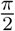. Additionally, the lower joints have a narrower distribution, up to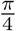. Among the *θ* angles, two DH *θ* angles (*θ*_1_ and *θ*_4_) are concentrated to near *−π* and *π*, while the other two (*θ*_2_ and *θ*_3_) have a wide distribution between *−π* and *π*. This observation indicates that the first joint, which demonstrates the rotation of the transmembrane region, and the fourth joint, the hip joint between the HR1 domain and the spike head, have a wider range of motion. This means that the main factor in the rotation of spike protein around the Z-axis is the transmembrane domain rotation. Also, the free rotation of the fourth (hip) joint allows the spike head to rotate freely and prime itself to bind to the human receptor. The restricted movement of the second and third joints (ankle and knee) might have different reasons, including the limited sampling during the simulations.

**Figure 2.**
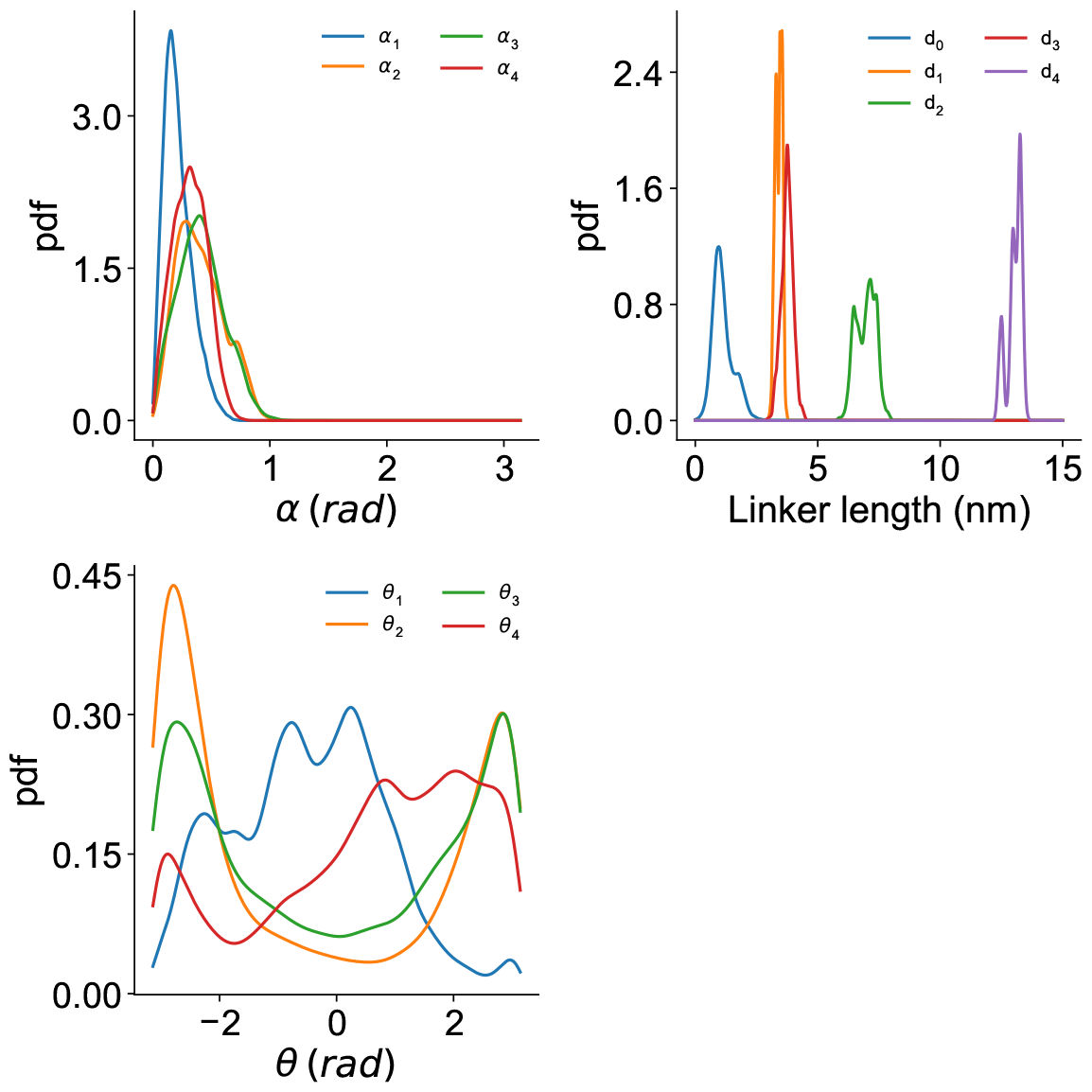
Distribution of Denavit-Hartenberg (DH) parameters in the simulations.

### 4.2 Euler angles and the SC parameters capture the global conformation

In the next step, we calculated the Euler angles and translational components (*α, β, γ, T*_*x*_, *T*_*y*_, *T*_*z*_) from the final DH transformation matrix. Also, we calculated the SC parameters (r, *θ*, and *φ*) for the tipping point. The distributions of the SC parameters and the Euler angles are shown in Fig. 3 and Fig.4, respectively. An interesting observation is that the Euler *α* and SC *φ* have a similar distribution, and the angles are up to 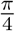. These two angles are functions of the DH *α* angles. The SC *φ* is simply the summation of the four DH *α* angles, while the Euler *α* is a more complex combination of the *α*s. One could expect that both the SC *φ* and Euler *α* can sample up to the sum of four DH *α*. But this is not simply the case, and these two global angles are restricted to far lower distribution (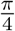 compared to the possible 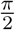). First, we might think it is because the spike can not bend more than a certain degree; otherwise, it would clash with the membrane. However, this is not the only reason, and the range is far narrower than clashing with the membrane. The first reason is that the final bending (Euler *α* or SC *φ*) is not a simple sum/combination of the DH *α*s, and the DH *θ*s also play a significant role in the spike conformation. Furthermore, it seems that the bending of the joints is not independent, and they are moving concertedly. This observation was previously proposed by Kapoor et al. [10]; however, they suggested that this is the effect of glycans. While it seems that this effect is, to some extent, the results of the glycan shield, it cannot be explained entirely based on the glycans, as we can see the correlated motion of the joints in some simulations of the naked spike. Figures 3 and 4 show that the Euler *γ* and SC *θ* angles span through the *−π* and *π* range. However, the distribution patterns are different, which is expected because the SC *θ* is the simple rotation of the tip point around the Z-axis. At the same time, the Euler *γ* is a combination of all joints’ conformations.

**Figure 3.**
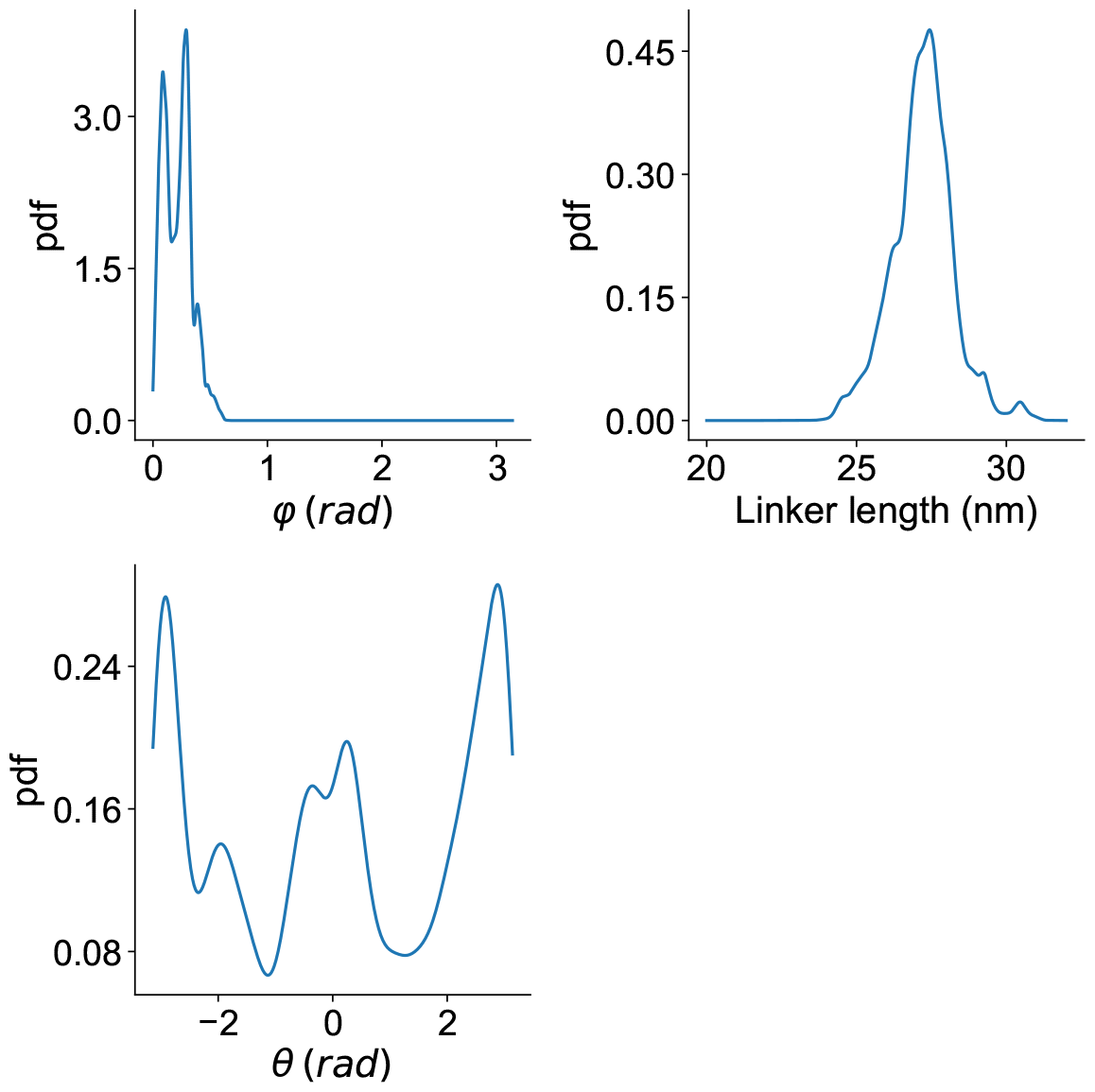
Distribution of spherical coordinates’ parameters from the simulations.

**Figure 4.**
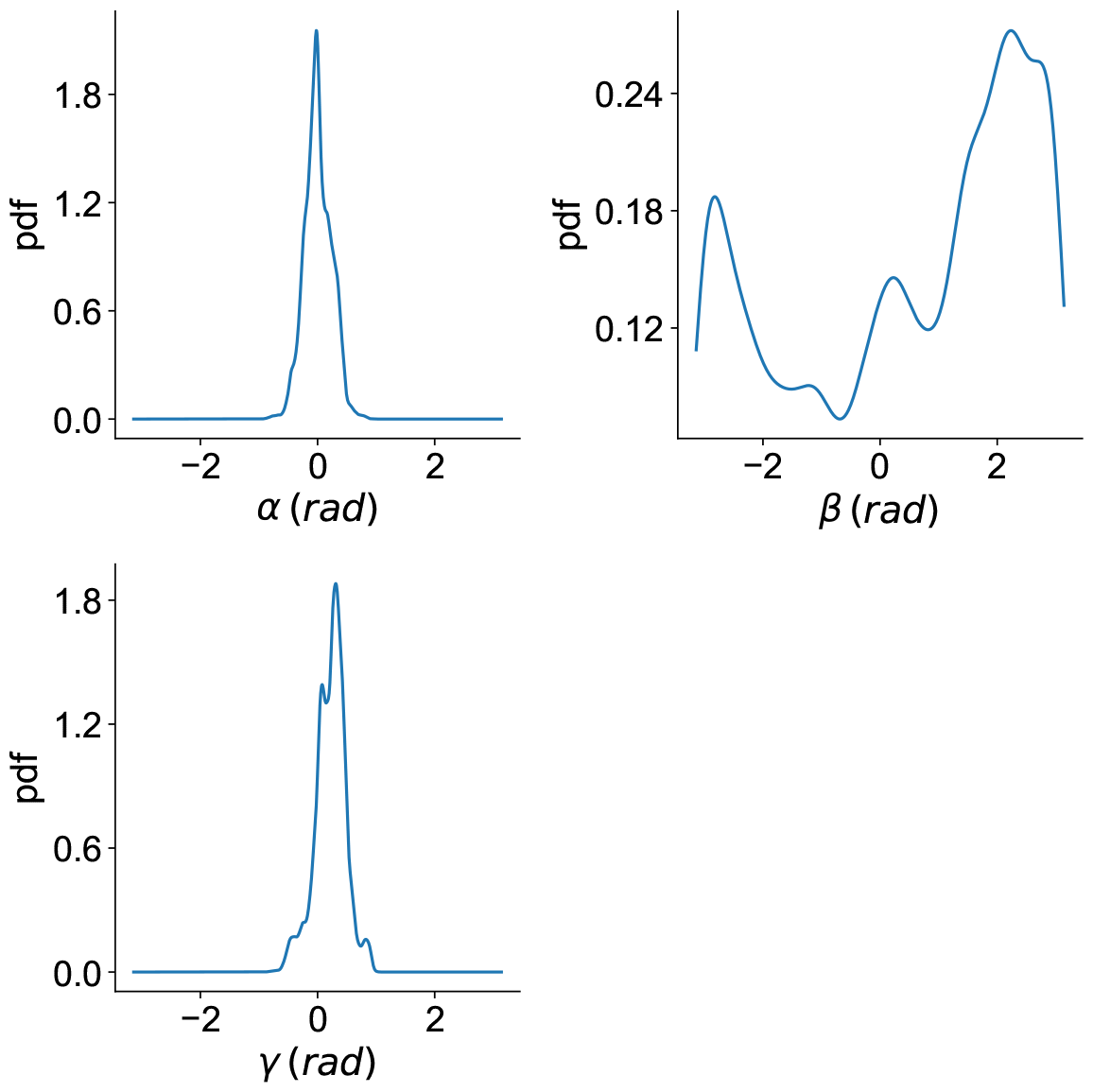
Distribution of DH-derived Euler angles from the simulations.

### 4.3 SC parameters cannot differentiate between two similar endpoints with different conformations

To look at the differences between the SC and DH parameters, we generate a set of DH parameters in which the linker lengths were fixed at the mean of the simulations (*d*_0_ = 1.74 *nm, d*_1_ = 3.45 *nm, d*_2_ = 7.00 *nm, d*_3_ = 3.94 *nm*, and *d*_4_ = 12.50 *nm*). Then, the *α* and *θ* angles were randomly assigned from their possible distribution (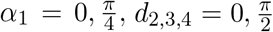). The parameter sets would be eliminated from the final data if, based on their predicted joint positions, they were clashing with the membrane.

This dataset shows a uniform distribution of possible conformation space for the DH and Euler parameters (Fig. S3 and S4). This robotic arm, the spike protein’s representatives, explores the extra-cellular space uniformly (Fig. S5). Among these generated robotic arm representations of the spike, we can find several instances in which the tip points are very close to each other, while the conformation of the spike is different. We identified similar instances among the conformations obtained from the actual simulations (Fig. 5). Based on the distance metric, we observed that the DH-derived transformation matrices correctly show the differences (the transformation distance metric is high). In contrast, the SC-derived transformation matrices are very close to each other (the distance metric is near zero) (Fig. S6). This difference is mainly because the DH rotation matrix is constructed based on all joints’ conformation and is reflected in the distance between two conformations. In contrast, the SC parameters only account for the tip points and do not include the joints’ conformation. This is also evident from the difference between the tip points’ distance and the RMSD between two robotic arms.

**Figure 5.**
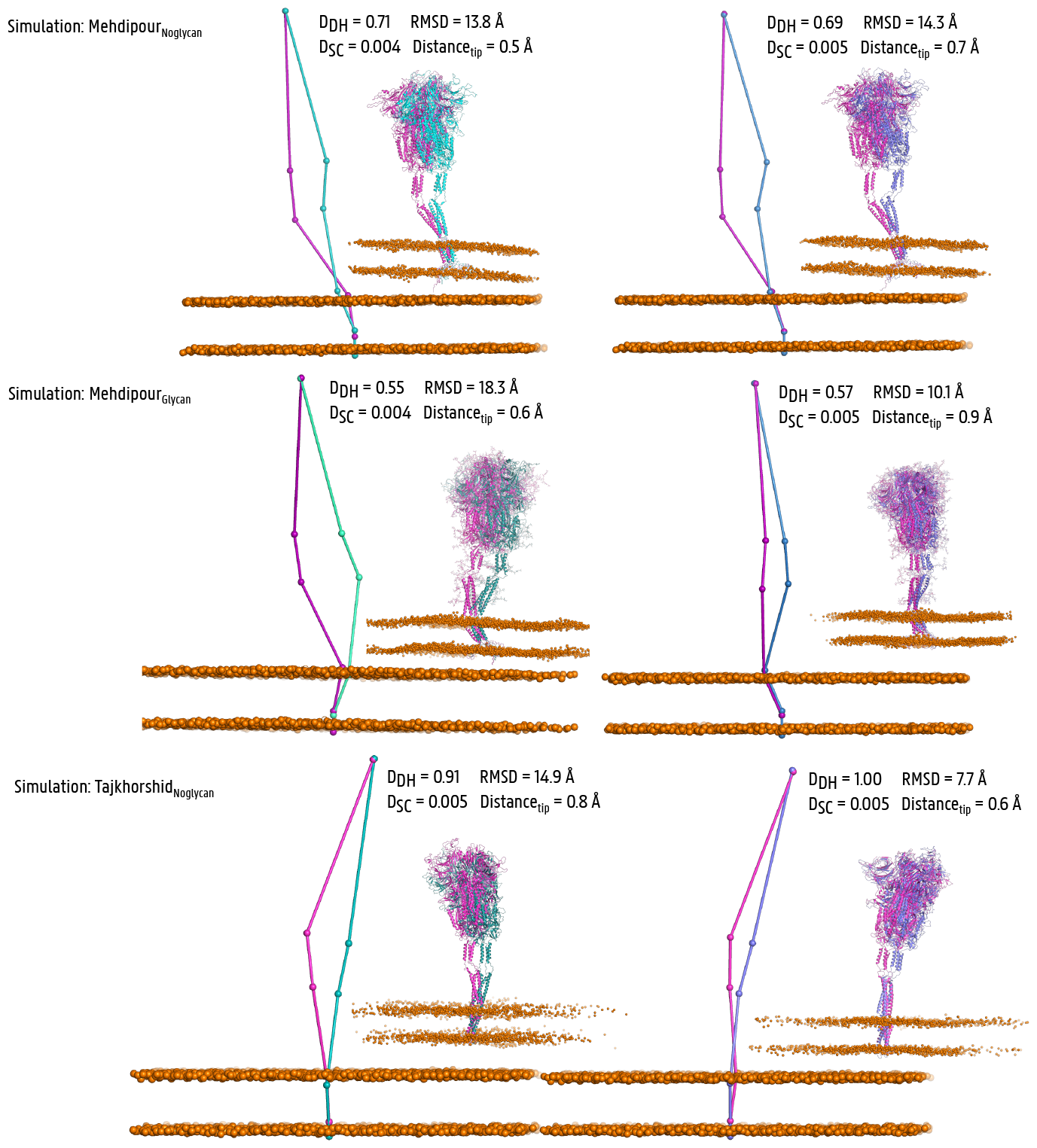
Instances of spike conformation with similar SC parameters and tip point position. *D*_*DH*_ and *D*_*SC*_ are the metric distances between two conformations based on DH-derived and SC-derived transformation matrices, respectively. RMSD is the root mean square distance between the points of the robotic arm representations of two conformations.

### 4.4 Rotational component magnifies upper joint change, translational component magnifies lower joint change

In different sets of simulations, we observed that the distribution of the transformation matrix shows some unique features that can help us understand the spike conformation space and the different conformational transitions. For example, in one of our glycosylated simulations, we see two clear conformation clusters based on the Euler angles. There is a transition between these clusters (Fig. 6). This transition is mainly based on the rotation of the tip (spike head) around the z-axis relative to the HR1 domain, i.e., changes in *θ*_4_, while *α*_2_ has a minor role on this transition. We can fit this transition by uniformly random changing of *θ*_4_, while *α*_2_ with other DH parameters kept near the mean values of this set.

**Figure 6.**
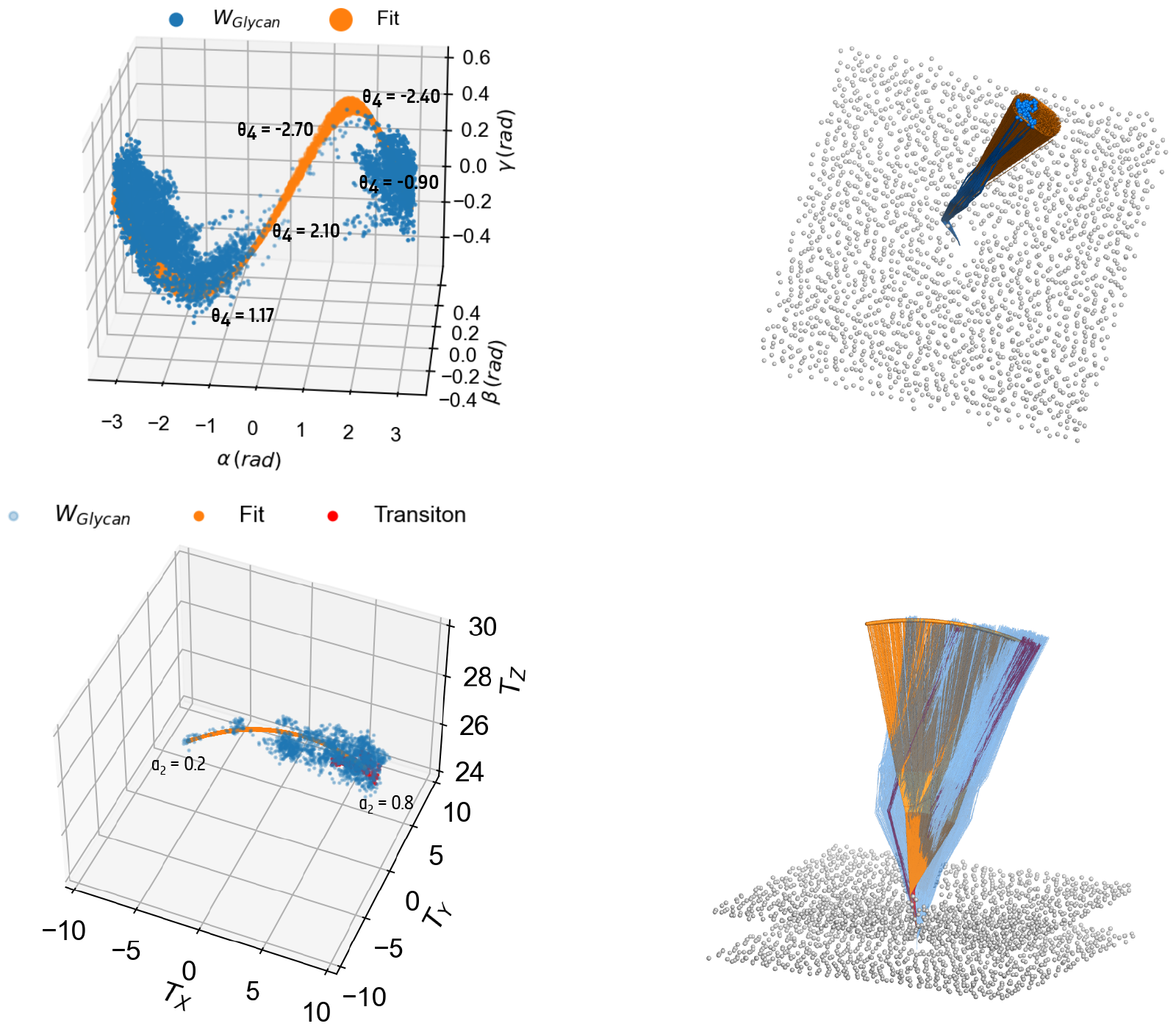
The Euler angles obtained from the rotational component of transformation matrices (top left panel) from one replicate of our fully glycosylated spike setup. The generated robotic arms are shown in the orange lines with randomly changing *θ*_4_ and *α*_2_ in the top and bottom panels, respectively. The red points in the lower panel are those in the transition part in the top panel.

However, the same transition has a negligible change in the translational component (Fig. 6). this is because the *α*_4_ is close to zero, and therefore, *θ*_4_ movement does not result in significant tip movement.

In contrast, if we look at the translational part, we observe a significant move away from the entirely straight conformation dominated by the change in *α*_2_. Interstingly, such considerable bending is not magnified in the rotation matrix because it is coming from a lower joint. It seems that the order of multiplication has an effect on the contribution of each joint to the final homogenous transformation.

## 5 Conclusion

Type 1 transmembrane receptors, such as the SARS-CoV-2 spike, are multi-domain flexible membrane proteins that play crucial roles in signaling and pathogen-host interactions. Their highly flexible nature makes it very challenging to gain detailed structural knowledge of these proteins in situ [3]. Here, combining MD simulations with a newly developed method inspired by forward kinematics approaches in robotics allows us to address this challenge and explore the conformation space of SARS-CoV-2 as a whole entity. Here, we performed MD simulations of the SARS-CoV-2 spike in the viral membrane. These simulations, plus the publicly available MD simulations of the spike, gave us a unique opportunity to explore the detailed definition of the conformational construct of the SARS-CoV-2 spike protein. Here, the DH parameters of the spike flexible hinges were used to build a homogeneous transformation matrix that identifies local and global changes in the overall conformational transitions.

In addition, this method can help us to sample the entire conformational space of the spike protein very fast, albeit in a very coarse-grained way. We can have uniformly sampled robotic arms representing the spike. In the next step, we aim to develop a workflow to convert the coarse-grained robotic arms to their atomistic models. These atomic models will be invaluable in several ways. First, seeds can be used for MD simulations to sample the local free energy surface. More importantly, as the structural biology of the cell receptors moves fast toward capturing the membrane proteins in their native environment using Cryo-ET, both the coarse-grained robotic arm and their converted atomic models can be used as a template for analyzing the conformational heterogeneity of the in-situ samples. Finally, having a few collective variables (three or six variables for the SC and DH approaches, respectively) will allow us to run enhanced sampling methods to sample the whole free energy surface of the spike effectively.

In conclusion, we established a new method to define the conformation of membrane receptors, which provides mechanistic insights. Our method provides a basis for sampling the conformational space of these receptors and will open the new possibility of looking at the conformational heterogeneity in situ.

## 6 Methods

### 6.1 Simulation protocol

The previously modeled spike protein system was used to generate the MD simulation inputs [20]. The transmembrane region of the spike protein was immersed in a membrane bilayer mimicking the viral membrane (25 % each of POPC and DOPC, 15 % POPI, 20 % POPE, and 5 % each of CHL1, CER160, and POPS) using CHARMM-GUI scripts [21]. Molecular dynamics simulations were carried out with GROMACS 2021 [22] and Charmm-36m forcefield parameters [23]. The spike and spike-glycan systems were solvated with the TIP3P water model, and the chargers were neutralized with 150 mM NaCl with the final size of the system 27*×*27*×*36 nm and a total of ^*∼*^2.528.000 atoms. Both the systems were energy minimized for 5,000 steepest-descent steps with all bonds with hydrogen atoms constrained using LINCS algorithm, followed by a six-step equilibration process with gradually reducing positional restraints applied on the heavy atoms and increasing the time step to 2 fs from an initial 1fs. The NVT equilibrations were performed using a Berendsen thermostat for 2.5 ns, and later, NPT equilibrations were performed using a Berendsen thermostat and a semi-isotropic barostat for 52 ns under periodic boundary conditions [24]. Long-range electrostatics were treated with Particle mesh Ewald summation with cubic interpolation and a cutoff of 0.12 nm [25]. The production simulations were performed using a Nose-Hoover thermostat [26] at 310 K and a semi-isotropic Parrinello-Rahman barostat [27] at 1 bar. Both glycosylated and non-glycosylated systems were simulated for 500 ns in triplicate. The MD trajectories were analyzed using Visual Molecular Dynamics (VMD) [28], MDAnalysis package [29], and in-house written codes.

### 6.2 RMSD of the robotic arms

The root-mean-square deviation (RMSD) between two spike robotic arms was calculated:

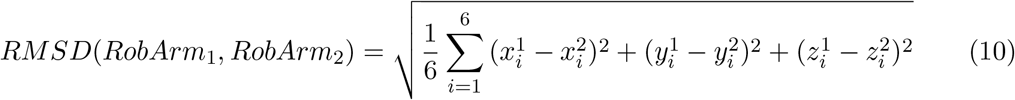

Where the summation is the distance between all six equivalent joints (base, four joints, and tip point) of two robotic arms.

## 7 Acknowledgments

The VSC (Flemish Supercomputer Center) and the EuroHPC supercomputer LUMI provided the high-performance computing resources and services used for performing the simulations in this work. A.R.M. acknowledges research support from Ghent University (BOF starting grant no. BOF.STG.2021.0037.01).

## A Appendix

### A.1 Details of Denavit-Hartenberg convention

In the definition of the Denavit-Hartenberg (DH) parameters (Fig. 2):

1. Each joint is placed on the origin of each frame (*A*_*i*_)
2. Z-axis (*Z*_*i*_) of each coordinate frame lies along the linker between the current joint (*A*_*i*_) and the subsequent joint (*A*_*i*+1_) in the chain.
3. X-axis (*X*_*i*_) of each coordinate frame will be perpendicular to the Z-axis of the current joint (*Z*_*i*_) and the previous joint (*Z*_*i−*1_) in the chain.
4. Then, by default, the Y-axis (*Y*_*i*_) of each coordinate frame will be defined as the axis perpendicular to both the Z-axis (*Z*_*i*_) and the X-axis (*X*_1_) of the frame.

For a robotic arm representation of a receptor protein, we can define the DH parameters for each joint as follows:

#### Joint i

*θ*_*i*_: Angle of rotation of *X*_*i−*1_ to *X*_*i*_ about *Z*_*i−*1_. The more intuitive definition of this angle is the dihedral angle between 4 consecutive joints (*A*_*i−*2_,*A*_*i−*1_, *A*_*i*_, *A*_*i*+1_)

*d*_*i*_: Distance between two consecutive joints (*A*_*i*_,*A*_*i*+1_).

*a*_*i*_: Distance along *X*_*i*_ from *A*_*i*_ (this parameter, in our case, is always 0, as the joints do not have a length.)

*α*_*i*_: Angle of rotation of *Z*_*i−*1_ to *Z*_*i*_ about *X*_*i*_. The more intuitive definition of this angle is the bond angle between 3 consecutive joints (*A*_*i−*1_, *A*_*i*_, *A*_*i*+1_)

For the spike protein, we have defined four joints, each with 3 degrees of freedom plus one translation from the base to the first joint; this means that we will have a nanorobotic arm with 13 degrees of freedom (13 DoF). DH parameters can be tabulated as:

We can write the homogeneous transformation matrix of the coordinates of the frame attached to joint i to the frame of joint i-1 based on three consecutive transformations:

1. Translation along (*Z*_*i*_) by *d*_*i*_
2. Rotation of (*Z*_*i*_) around (*X*_*i*_) by *α*_*i*_
3. Rotation of (*X*_*i*_) around (*Z*_*i−*1_) by *θ*_*i*_

#### Translation along Z-axis by *d*_*i*_

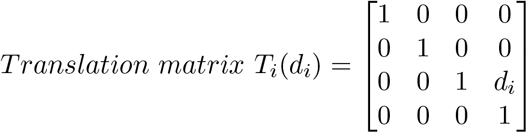

#### Rotation of Z-axis by *α*_*i*_

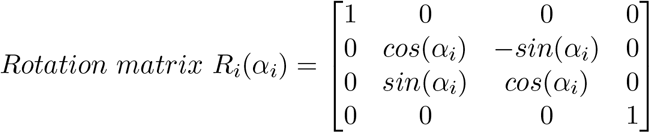

#### Rotation of X-axis by *θ*_*i*_

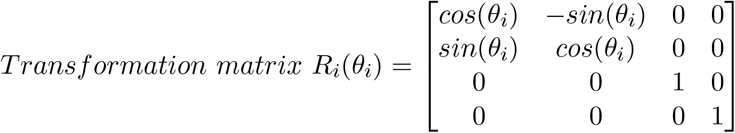

By multiplying these rotation and translation matrices, we will get the final transformation matrix for the transformation from i to i+1.

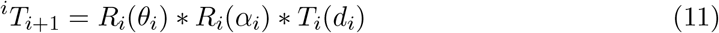

#### Transformation matrix from joint i+1 to i (^*i*^*T*_*i*+1_)

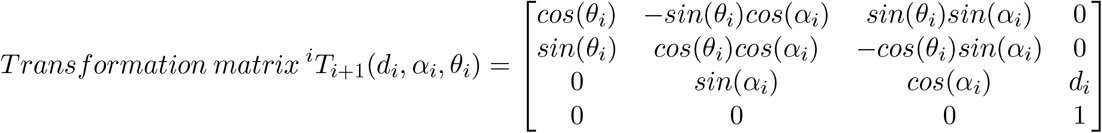

The only exception is the transformation from the base to joint one (*A*_1_), which is a translation-only operation:

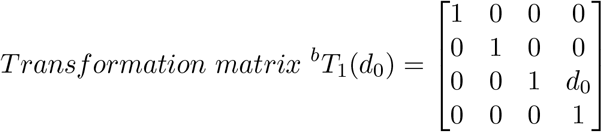

Then, the transformation matrix of the whole chain to have the coordinates of the endpoint effector (tip of the S) in the base coordinate system will be:

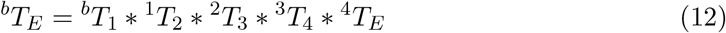

The final transformation matrix can b e written as:

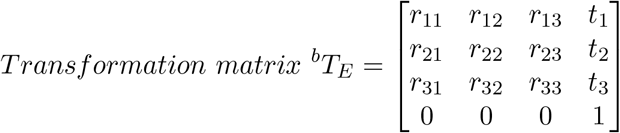

Each element of the transformation matrix is composed of:

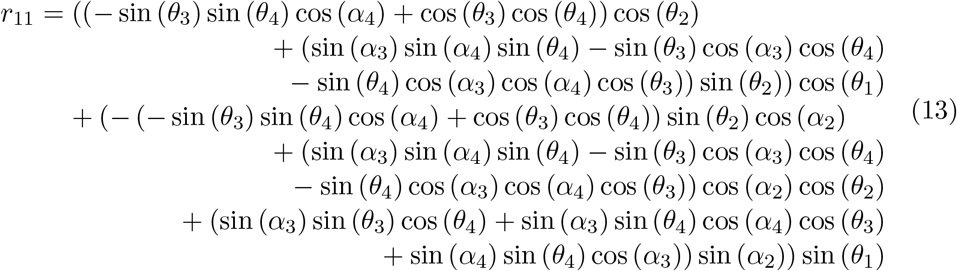

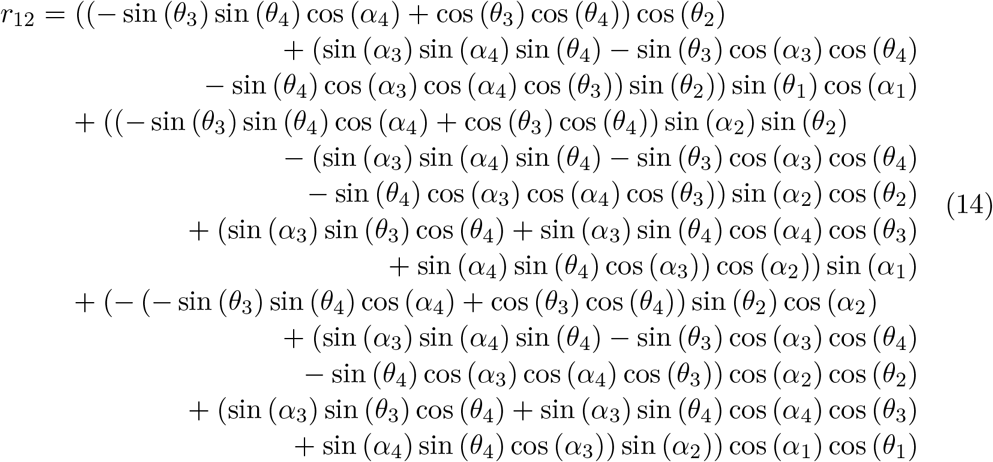

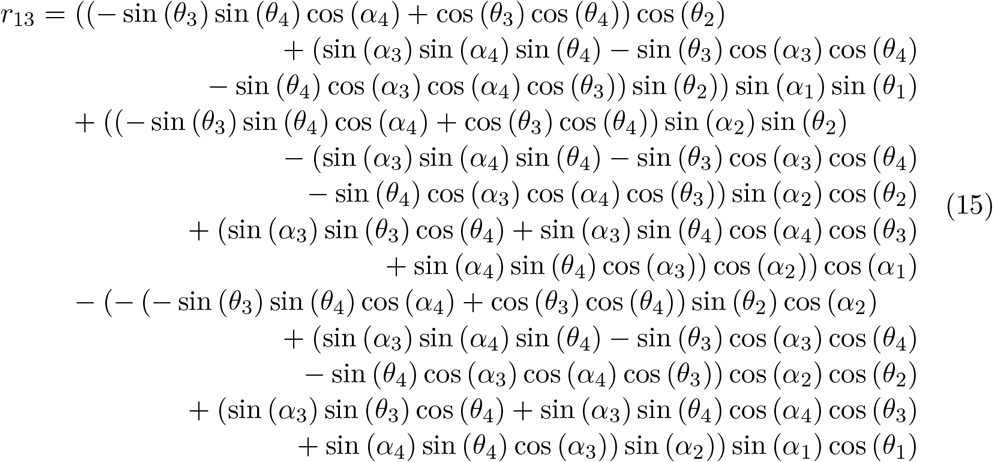

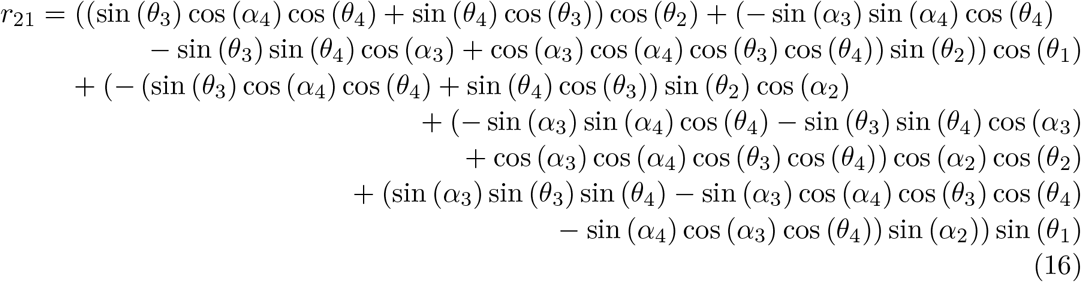

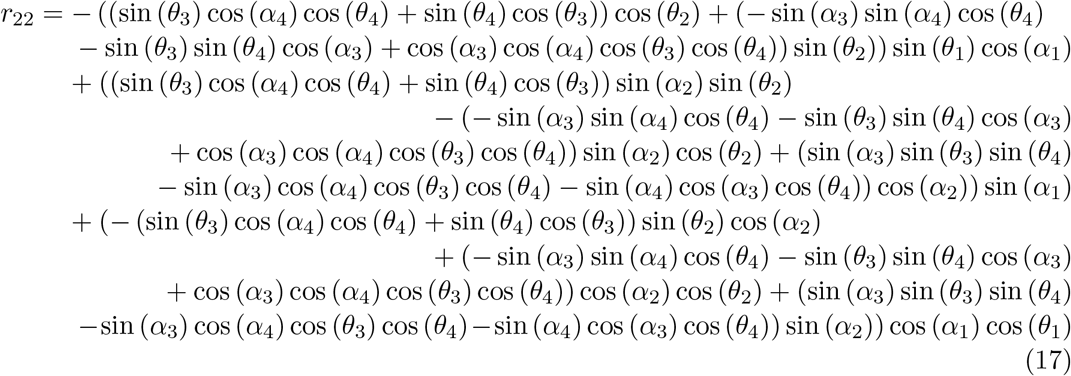

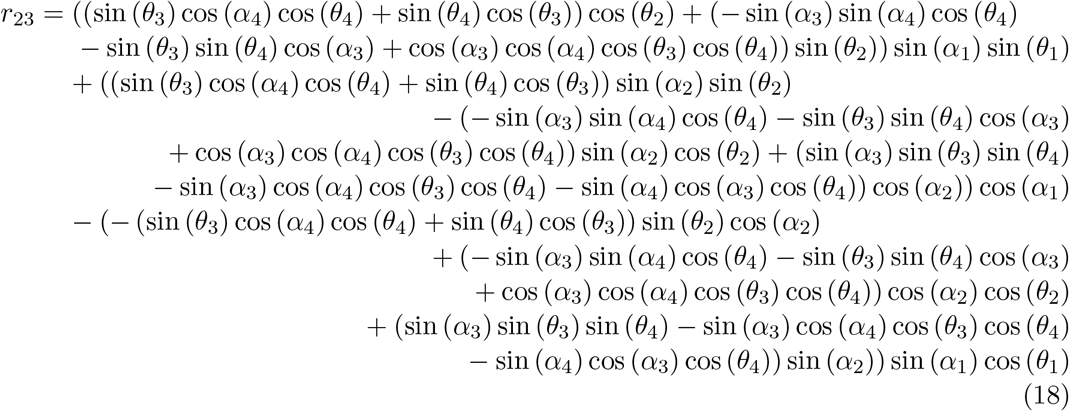

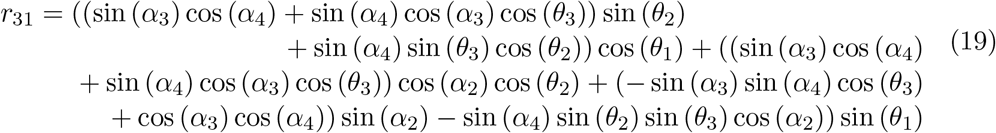

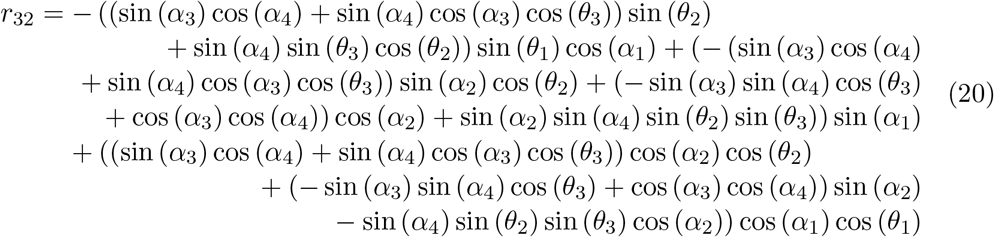

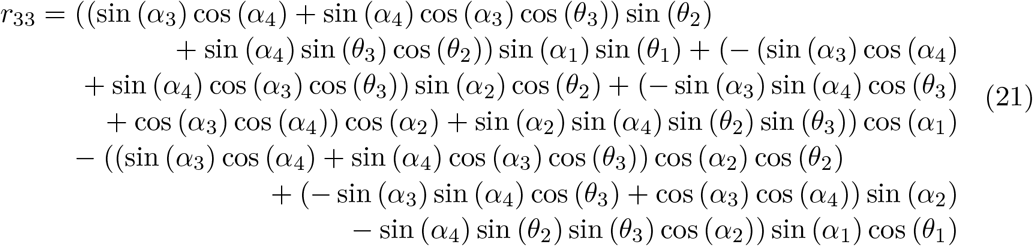

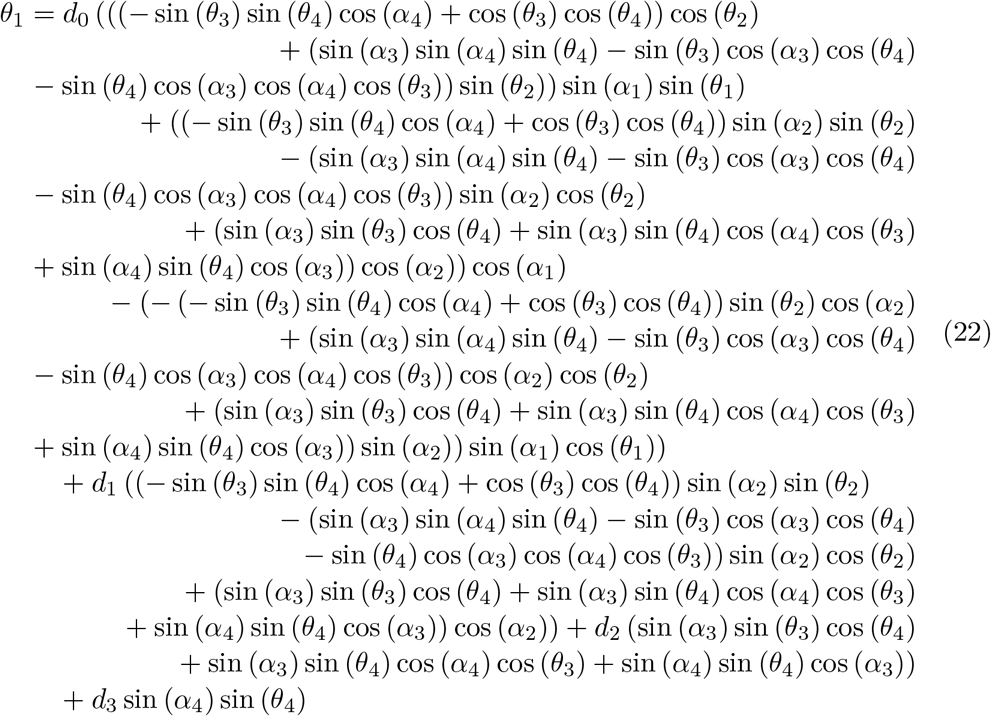

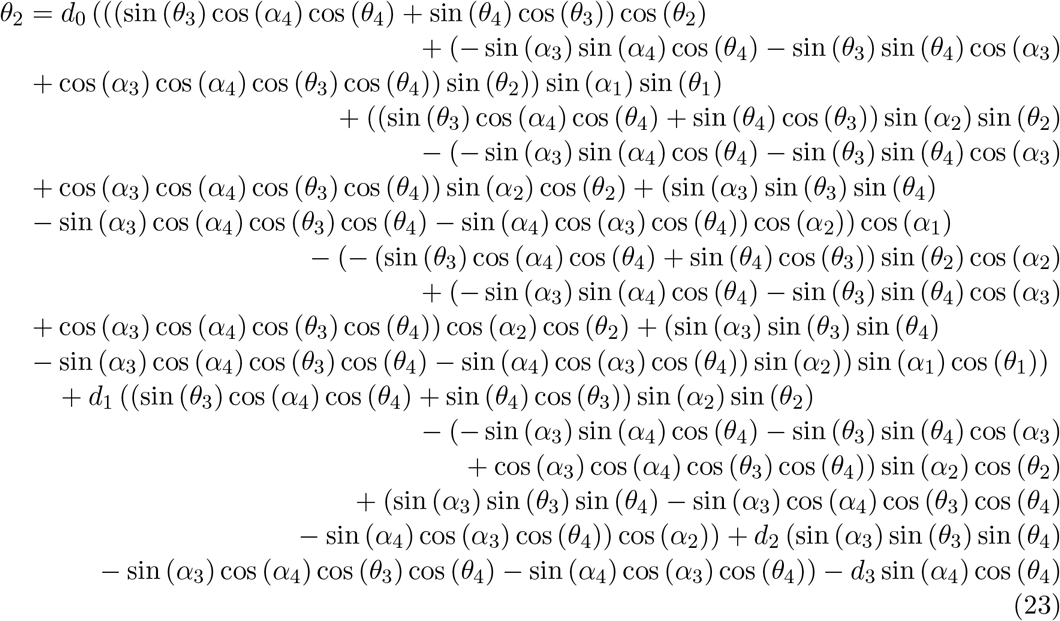

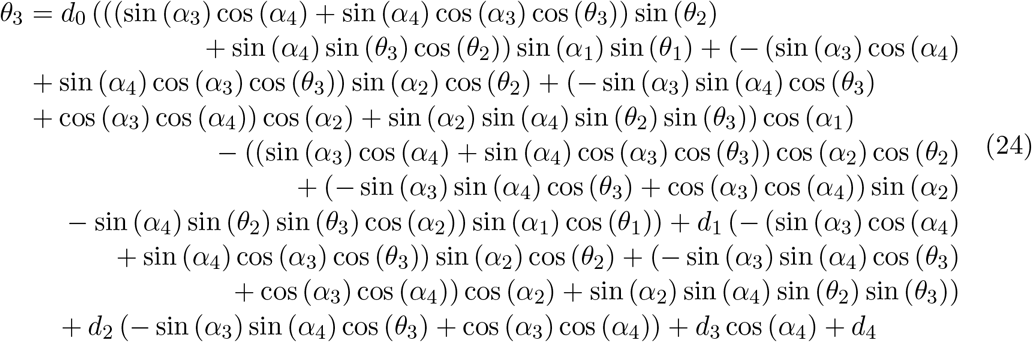

## Notes

### Competing Interest Statement

The authors have declared no competing interest.

## References

[1] Lucas M. P. Chataigner, Nadia Leloup, Bert J. C. Janssen (2020) Structural Perspectives on Extracellular Recognition and Conformational Changes of Several Type-I Transmembrane Receptors. Front. Mol. Biosci., 7, 129.

[2] Katrine Bugge, Kresten Lindorff-Larsen, Birthe B Kragelund (2016) Understanding single-pass transmembrane receptor signaling from a structural viewpoint-what are we missing? FEBS J., 283, 4424–4451.

[3] Kai Cai, Xuewu Zhang, Xiao-chen Bai (2022) Cryo-electron Microscopic Analysis of Single-Pass Transmembrane Receptors. Chem. Rev. 122, 13952–13988.

[4] João P. Luís, Ana I. Mata, Carlos J. V. Simões, Rui M. M. Brito (2022) Conformational Dynamics of the Soluble and Membrane-Bound Forms of Interleukin-1 Receptor Type-1: Insights into Linker Flexibility and Domain Orientation. Int. J. Mol. Sci., 23, 2599.

[5] Chao-Yie Yang (2020) Comparative Analyses of the Conformational Dynamics Between the Soluble and Membrane-Bound Cytokine Receptors. Sci Rep 10, 7399.

[6] Bryant Gipson, David Hsu, Lydia E Kavraki, Jean-Claude Latombe (2012) Computational models of protein kinematics and dynamics: beyond simulation. Annu. Rev. Anal. Chem., 5, 273–291.

[7] Turoňová, Beata, Mateusz Sikora, Christoph Schuürmann, Wim JH Hagen, Sonja Welsch, Florian EC Blanc, Söoren von Buülow, Michael Gecht, Katrin Bagola, Cindy Höorner, Ger van Zandbergen,bShyamal Mosalaganti, Andre Schwarz, Roberto Covino, Michael D. Muühlebach, Gerhard Hummer, Jacomine Krijnse Locker, Martin Beck (2020) In situ structural analysis of SARS-CoV-2 spike reveals flexibility mediated by three hinges. Science, 370, 203–208.

[8] Ahmad Reza Mehdipour, Gerhard Hummer (2021) Dual nature of human ACE2 glycosylation in binding to SARS-CoV-2 spike. PNAS, 118, e2100425118.

[9] Lorenzo Casalino, Zied Gaieb, Jory A. Goldsmith, Christy K. Hjorth, Abigail C. Dommer, Aoife M. Harbison, Carl A. Fogarty, Emilia P. Barros, Bryn C. Taylor, Jason S. McLellan, Elisa Fadda, Rommie E. Amaro (2020) Beyond Shielding: The Roles of Glycans in the SARS-CoV-2 Spike Protein. ACS Cent. Sci., 6, 1722–1734.

[10] Karan Kapoor, Tianle Chen, Emad Tajkhorshid (2022) Posttranslational modifications optimize the ability of SARS-CoV-2 spike for effective interaction with host cell receptors. PNAS, 119, e119761119.

[11] David Chmielewski, Eric A Wilson, Grigore Pintilie, Peng Zhao, Muyuan Chen, Michael F Schmid, Graham Simmons, Lance Wells, Jing Jin, Abhishek Singharoy, Wah Chiu (2023) Structural insights into the modulation of coronavirus spike tilting and infectivity by hinge glycans. Nat. Commun., 14, 7175.

[12] Yeol Kyo Choi, Yiwei Cao, Martin Frank, Hyeonuk Woo, Sang-Jun Park, Min Sun Yeom, Tristan I. Croll, Chaok Seok, Wonpil Im (2021) Structure, Dynamics, Receptor Binding, and Antibody Binding of the Fully Glycosylated Full-Length SARS-CoV-2 Spike Protein in a Viral Membrane. J. Chem. Theory Comput. 17, 4, 2479–2487

[13] Dinesh Manocha, Yunshan Zhu, William Wright (1995) Conformational analysis of molecular chains using nano-kinematics. CABIOS, 11, 71–85.

[14] Hernan Stamati, Amarda Shehu, Lydia Kavraki (2007) Computing Forward Kinematics for Protein-like linear systems using Denavit-Hartenberg Local Frames. https://www.researchgate.net/publication/251264464?Computing?Forward?Kinematics?for?Protein-like?linear?systems?using?Denavit-Hartenberg?Local?Frames.

[15] J. Denavit, R. S. Hartenberg (1955) A Kinematic Notation for Lower-Pair Mechanisms Based on Matrices. J. Appl. Mech. 22, 215–221.

[16] Raffaele Di Gregorio (2024) Metrics proposed for measuring the distance between two rigid-body poses: review, comparison, and combination. Robotica. 42, 302–318.

[17] Mariona Torrens-Fontanals, Alejandro Peralta-García, Carmine Talarico, Ramon Guixà-González, Toni Giorgino, Jana Selent (2021) SCoV2-MD: a database for the dynamics of the SARS-CoV-2 proteome and variant impact predictions. Nucl. Acids Res., 50, D858–D866.

[18] Hyeonuk Woo, Sang-Jun Park, Yeol Kyo Choi, Taeyong Park, Maham Tanveer, Yiwei Cao, Nathan R Kern, Jumin Lee, Min Sun Yeom, Tristan I Croll, Chaok Seok, Wonpil Im (2020) Developing a Fully Glycosylated Full-Length SARS-CoV-2 Spike Protein Model in a Viral Membrane. J. Phys. Chem. B, 124, 7128–7137.

[19] Mahdi Ghorbani, Bernard R. Brooks, Jeffery B. Klauda (2021) Exploring dynamics and network analysis of spike glycoprotein of SARS-COV-2. Biophys. J., 120, 2902–2913.

[20] Mateusz Sikora, Söoren von Buülow, Florian E. C. Blanc, Michael Gecht, Roberto Covino, Gerhard Hummer (2021) Computational epitope map of SARS-CoV-2 spike protein. PLoS Comput. Biol., 17, e1008790.

[21] Emilia L. Wu, Xi Cheng, Sunhwan Jo, Huan Rui, Kevin C. Song, Eder M. Dávila-Contreras, Yifei Qi, Jumin Lee, Viviana Monje-Galvan, Richard M. Venable, Jeffery B. Klauda, Wonpil Im (2014) CHARMM-GUI Membrane Builder toward realistic biological membrane simulations. J. Comput. Chem., 35, 1997–2004.

[22] Mark James Abraham, Teemu Murtola, Roland Schulz, Szilárd Páll, Jeremy C. Smith, Berk Hess, Erik Lindahl (2015) GROMACS: High performance molecular simulations through multi-level parallelism from laptops to supercomputers. SoftwareX, 1-2, 19–25.

[23] Robert B. Best, Xiao Zhu, Jihyun Shim, Pedro E. M. Lopes, Jeetain Mittal, Michael Feig, and Alexander D. MacKerell, Jr (2012) Optimization of the Additive CHARMM All-Atom Protein Force Field Targeting Improved Sampling of the Backbone ϕ, Ψ and Side-Chain χ1 and χ2 Dihedral Angles. J. Chem. Theo. Comput., 8, 3257–3273.

[24] H. J. C. Berendsen; J. P. M. Postma; W. F. van Gunsteren; A. DiNola; J. R. Haak (1984) Molecular-Dynamics with Coupling to an External Bath. J. Chem. Phys., 81, 3684–3690.

[25] Tom Darden; Darrin York; Lee Pedersen (1993) Particle mesh Ewald: An N.log(N) method for Ewald sums in large systems. J. Chem. Phys., 98, 10089–10092.

[26] William G. Hoover (1985) Canonical dynamics: Equilibrium phase-space distributions. Phys. Rev. A, 31, 1695–1697.

[27] Michelle Parrinello, A. Rahman (1981) Polymorphic Transitions in Single-Crystals - a New Molecular-Dynamics Method. J. Appl. Phys., 52, 7182–7190.

[28] William Humphrey, Andrew Dalke, Klaus Schulten (1996) VMD: visual molecular dynamics. J. Mol. Graph., 14, 27–38.

[29] Michaud-Agrawal, N., Denning, E. J., Woolf, T. B. and Beckstein, O. (2012) MDAnalysis: a toolkit for the analysis of molecular dynamics simulations. J. Comput. Chem., 32, 2319–2327.

