## Supplementary figures and tables for "Robotic-inspired approach to multi-domain membrane receptor conformation space: theory and SARS-CoV-2 spike protein case study"

Table 1: Points definition

| **Point** | **Definition** |
|---|---|
| Base | Z coordinate of $COM^a$ of P atoms of lower leaflet lipid molecules + XY coordinates of Joint 1 |
| Joint 1 | COM of $C_\alpha$ atoms of residues 1233-1237 of three chains A B C |
| Joint 2 | COM of $C_\alpha$ atoms of residues 1207-1215 of three chains A B C |
| Joint 3 | COM of $C_\alpha$ atoms of residues 1160-1174 of three chains A B C |
| Joint 4 | COM of $C_\alpha$ atoms of residues 11141-1146 of three chains A B C |
| End Effector | COM of $C_\alpha$ atoms of residues 332-442 of three chains A B C |

[a] COM Center of mass

Table 2: Simulation data used in this study

| **Source** | **Length** | **Description** | **Code name** |
|---|---|---|---|
| Mehdipour lab | 3 × 0.5 $\mu$s | Full length glycosylated spike | $Mehdipour_{Glyc}$ |
| Mehdipour lab | 3 × 0.5 $\mu$s | Full length nonglycosylated spike | $Mehdipour_{Noglyc}$ |
| Hummer lab | 2.5 $\mu$s | Four copies of Full length glycosylated spike | $Hummer_{Glyc}$ |
| Tajkhorshid lab | 5 $\mu$s | Full length glycosylated spike | $Tajkhorshid_{Glyc}$ |
| Tajkhorshid lab | 5 $\mu$s | Full length nonglycosylated spike | $Tajkhorshid_{Noglyc}$ |
| Im lab | 16 × 1.28 $\mu$s | Full length glycosylated spike | $Im_{Glyc}$ |
| Klauda lab | 1 $\mu$s | Full length glycosylated spike | $Klauda_{glyc}$ |

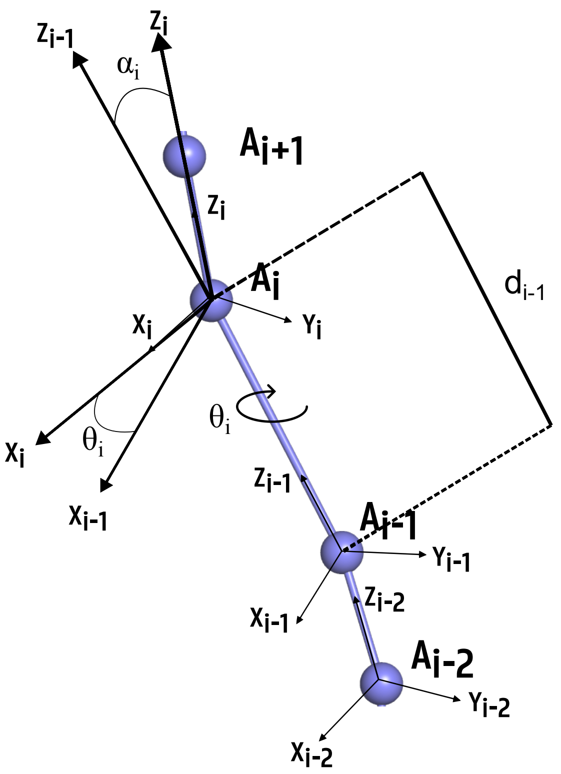


Figure S 1: **Denavit-Hartenberg (DH) convention and parameters**.

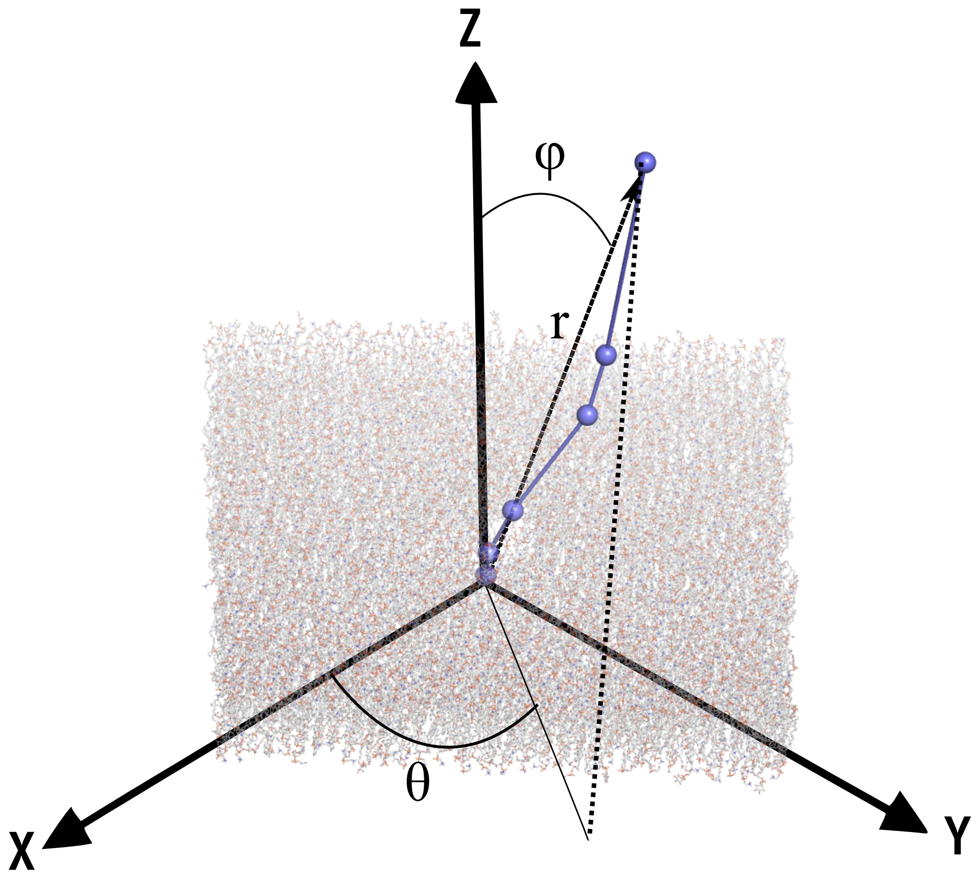


Figure S 2: **Spherical coordinates convention and parameters**.

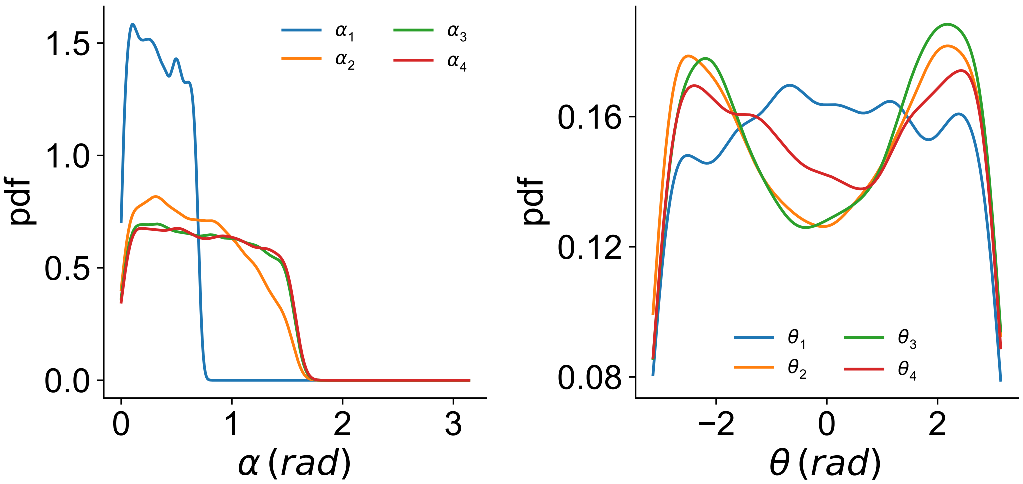


Figure S 3: **Distribution of Denavit-Hartenberg (DH) parameters for the generated robotic arm of spike**.

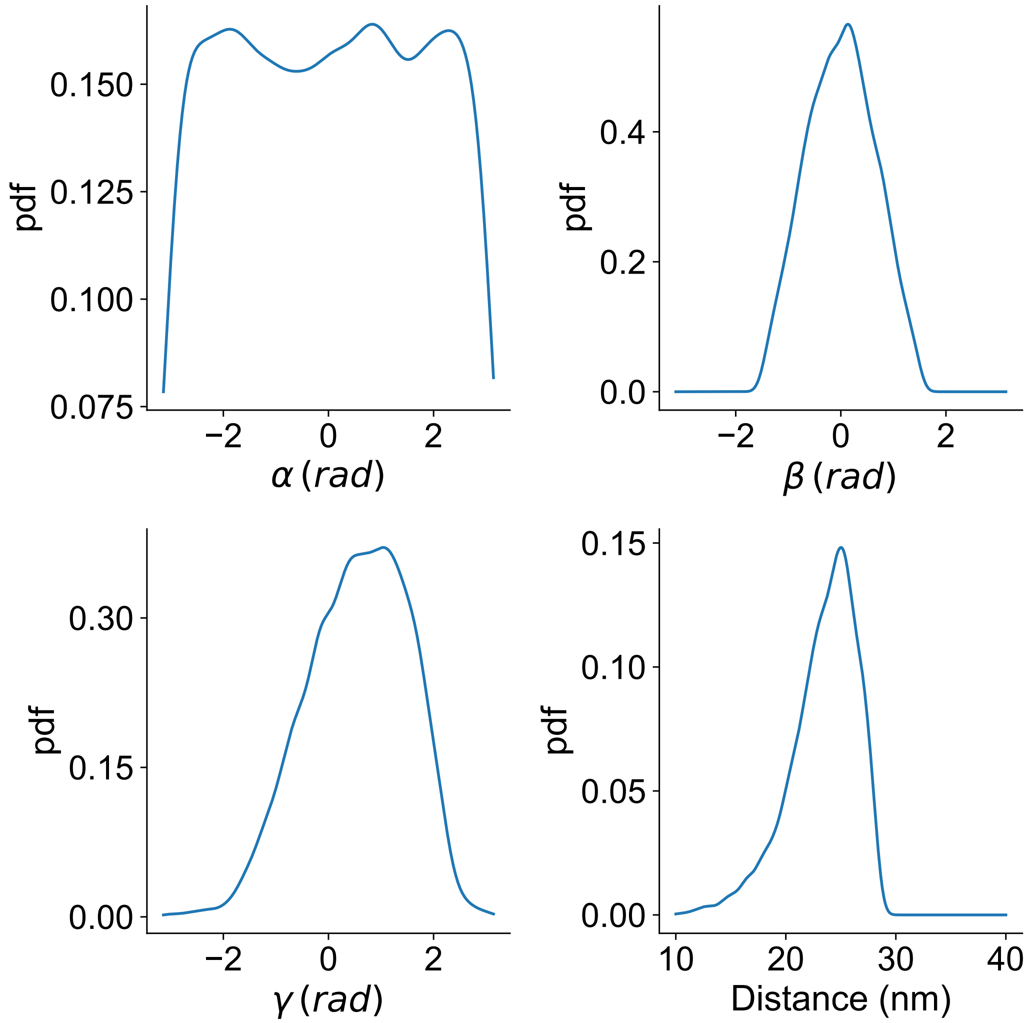


Figure S 4: **Spherical coordinate parameters for the generated robotic arm of spike**.

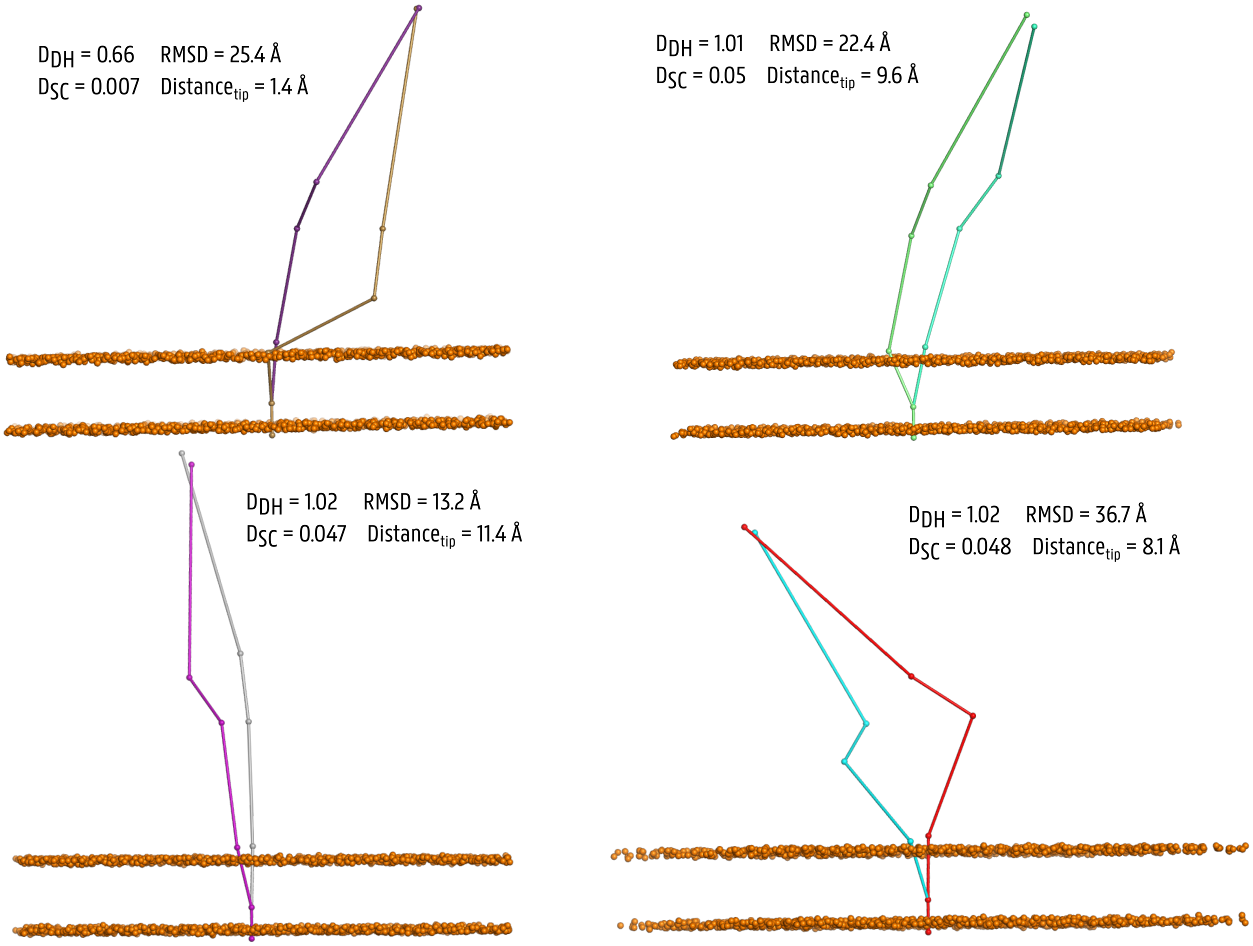


Figure S 5: **Generated robotic arms with similar SC parameters and tip point position**.
